# Sodium regulates PLC and IP_3_R-mediated calcium signalling in invasive breast cancer cells

**DOI:** 10.1101/2022.08.10.503447

**Authors:** Andrew D. James, Katherine P. Unthank, Isobel Jones, Nattanan Sajjaboontawee, Rebecca E. Sizer, Sangeeta Chawla, Gareth J.O. Evans, William J. Brackenbury

## Abstract

Intracellular Ca^2+^ signalling and Na^+^ homeostasis are inextricably linked via ion channels and co-transporters, with alterations in the concentration of one ion having profound effects on the other. Evidence indicates that intracellular Na^+^ concentration ([Na^+^]_i_) is elevated in breast tumours, and that aberrant Ca^2+^ signalling regulates numerous key cancer hallmark processes. The present study therefore aimed to determine the effects of Na^+^ depletion on intracellular Ca^2+^ handling in metastatic breast cancer cell lines. The relationship between Na^+^ and Ca^2+^ was probed using fura-2 and SBFI fluorescence imaging and replacement of extracellular Na^+^ with equimolar N-methyl-D-glucamine (0Na^+^/NMDG) or choline chloride (0Na^+^/ChoCl). In triple-negative MDA-MB-231 and MDA-MB-468 cells and Her2+ SKBR3 cells, but not ER+ MCF-7 cells, 0Na^+^/NMDG and 0Na^+^/ChoCl resulted in a slow, sustained depletion in [Na^+^]_i_ that was accompanied by a rapid and sustained increase in intracellular Ca^2+^ concentration ([Ca^2+^]_i_). Application of La^3+^ in nominal Ca^2+^-free conditions had no effect on this response, ruling out reverse-mode NCX activity and Ca^2+^ entry channels. Moreover, the Na^+^-linked [Ca^2+^]_i_ increase was independent of membrane potential hyperpolarisation (NS-1619), but was inhibited by pharmacological blockade of IP_3_ receptors (2-APB), phospholipase C (PLC, U73122) or following depletion of endoplasmic reticulum Ca^2+^ stores (cyclopiazonic acid). Thus, Na^+^ is linked to PLC/IP_3_-mediated activation of endoplasmic reticulum Ca^2+^ release in metastatic breast cancer cells and this may have an important role in breast tumours where [Na^+^]_i_ is perturbed.

## Introduction

Early detection has played an important role for timely clinical intervention in breast cancer; however, development of novel therapies which selectively kill malignant cells remains a central research goal. Key hallmarks of cancer include unlimited replicative capacity, self-sufficiency of growth, apoptosis resistance, angiogenesis and ultimately tissue invasion (Hanahan & Weinberg, 2016). Importantly, intracellular ion signalling pathways (in particular Ca^2+^ and Na^+^ signalling) play a key role in regulating many of these hallmark processes in breast cancer (Leslie *et al*., 2019; Bruce & James, 2020; Capatina *et al*., 2022; Yang & Brackenbury, 2022), and thus ion signalling may be a novel therapeutic locus.

Intracellular [Na^+^] ([Na^+^]_i_) and intracellular [Ca^2+^] ([Ca^2+^]_i_) signalling is achieved via a multiplicity of ion channels and transporters located both at the plasma membrane and on intracellular organelles. These channels harness the electrochemical gradient of Na^+^ and Ca^2+^ across cell membranes to achieve their function, with [Na^+^]_i_ and [Ca^2+^]_i_ typically tightly regulated by various ion transporters (Bruce, 2018; Leslie *et al*., 2019). Importantly, Ca^2+^ and Na^+^ signalling are inextricably linked via mechanisms mutually regulated by both ions, such as the plasma membrane Na^+^/Ca^2+^ exchanger (NCX; which is responsible for bulk export of Ca^2+^ from the cytosol) and the mitochondrial Na^+^/Ca^2+^/Li^+^ exchanger (which regulates mitochondrial Ca^2+^ and Na^+^ and thus bioenergetics (Hernansanz-Agustín *et al*., 2020)). Thus, changes in how the cell handles one ion would be expected to impact upon the other. Moreover, the interplay between [Na^+^]_i_ and [Ca^2+^]_i_ handling has important implications in breast cancer since [Na^+^]_i_ is elevated in malignant breast cancer cells, particularly in those exhibiting aberrant Na^+^ channel expression (Cameron *et al*., 1980; Ouwerkerk *et al*., 2007; Yang *et al*., 2020; James *et al*., 2022).

Emerging evidence implicates altered Na^+^ and Ca^2+^ signalling in the progression of breast cancer (Leslie *et al*., 2019; Bruce & James, 2020). For example, aberrantly expressed voltage-gated Na^+^ channels (VGSCs) drive metastasis (Fraser *et al*., 2005; Brackenbury *et al*., 2007; Brisson *et al*., 2013; Driffort *et al*., 2014; Nelson *et al*., 2014, 2015b, 2015a; Bon *et al*., 2016), elevations in intracellular Na^+^ correlate with malignancy and treatment response (Ouwerkerk *et al*., 2007; James *et al*., 2022) and upregulation of numerous Ca^2+^ channels and ATPases (and thus alterations in Ca^2+^ handling) are linked with breast cancer progression and poor prognosis (VanHouten *et al*., 2010; Feng *et al*., 2010; Middelbeek *et al*., 2012; Davis *et al*., 2014; Jeong *et al*., 2016; So *et al*., 2019). The apparent elevation in [Na^+^]_i_ exhibited by breast cancer cells might be expected to affect Na^+^-dependent Ca^2+^ handling mechanisms; for example, NCX, which can operate in reverse, Ca^2+^ entry mode following changes in [Na^+^]_i_ (Pappalardo *et al*., 2014; Verkhratsky *et al*., 2018). Nevertheless, the interplay between Na^+^ and Ca^2+^ handling in the context of cancer is poorly understood (Leslie *et al*., 2019; Bruce & James, 2020).

The present study aimed to determine the effects of Na^+^ depletion on intracellular Ca^2+^ handling in metastatic breast cancer cell lines in order to probe the relationship between Na^+^ and Ca^2+^ in breast cancer. A commonly recognised method to probe the role of Na^+^ transport in cell physiology is by replacement of extracellular Na^+^ with equimolar N-methyl-D-glucamine (0Na/NMDG) or choline chloride (0Na/ChoCl). These replacement cations maintain the osmotic balance across the cell membrane, yet are not transported via Na^+^ channels or transporters. Breast cancer cells can be subdivided into three main categories based on expression of the estrogen receptor (ER), progesterone receptor and human epidermal growth factor 2 (Her2). In triple-negative MDA-MB-231 and MDA-MB-468 cells (lacking all three receptors) and Her2+ SKBR3 cells, removal of Na^+^ from the extracellular space led to a slow, sustained depletion in [Na^+^]_i_ that was accompanied by a rapid and sustained increase in intracellular [Ca^2+^]_i_. Interestingly, this response was absent in ER+ MCF-7 cells. The observed [Ca^2+^]_i_ transient was not due to Ca^2+^ entry from the extracellular space, ruling out reverse-mode NCX activity. Moreover, it was independent of membrane potential (V_m_) and was inhibited by depletion of the endoplasmic reticulum Ca^2+^ store or by pharmacological blockade of inositol (1,4,5) trisphosphate (IP_3_) receptors (IP_3_Rs) or phospholipase C (PLC). These data reveal a previously unreported Na^+^-linked activation of endoplasmic reticulum Ca^2+^ release in metastatic breast cancer cells, which may have an important role in breast tumours where [Na^+^]_i_ is elevated or perturbed. Moreover, the dramatic Ca^2+^ release observed upon Na^+^ depletion serves as a cautionary note for those utilising similar Na^+^ replacement methods to study the role of Na^+^ transport in cellular processes.

## Materials and methods

### Cell Culture

Cells were cultured at 37 °C in a humidified atmosphere of air/CO_2_ (95:5%) in Dulbecco’s modified Eagle’s medium (DMEM 219690-35, Thermo Fisher Scientific) supplemented with 4 mM L-glutamine and 5% foetal bovine serum, as described previously (Ding & Djamgoz, 2004; Pan & Djamgoz, 2008; Yang *et al*., 2012; Brisson *et al*., 2013). Cells were routinely tested for mycoplasma using the DAPI method (Uphoff *et al*., 1992). MDA-MB-231 and MCF-7 cells were gifts from M. Djamgoz (Imperial College London) and SKBR3 cells were a gift from J. Rae (University of Michigan). Molecular identity of MDA-MB-231, MCF-7 and SKBR3 cells was verified by short tandem repeat analysis (Masters et al. 2001). MDA-MB-468 cells were obtained for this study directly from the American Type Culture Collection (ATCC).

### Fura-2 and SBFI fluorescence microscopy

Cells were seeded onto glass coverslips and allowed to adhere overnight. Cells were loaded with either fura-2 AM (4 μM) or SBFI AM (4 μM) in HEPES-buffered physiological saline solution (HEPES-PSS: 144 mM NaCl, 5.4 mM KCl, 1 mM MgCl_2_, 1.28 mM CaCl_2_, 5 mM HEPES and 5.6 mM glucose, pH 7.2) with 0.08% Pluronic F-127 at room temperature (fura-2 AM, 40 min; SBFI AM, 2 hours). Cells were then rinsed once with HEPES-PSS and incubated in dye-free HEPES-PSS to allow uncleaved dye to re-equilibrate (fura-2 AM, 20 min; SBFI AM, 40 min). For all experiments except those shown in Supplementary Figure 1, loaded coverslips were mounted within a perfusion chamber (RC26G, Warner Instruments) fitted to an imaging system comprising an Eclipse TE-200 inverted microscope (Nikon), a Plan Fluor ELWD 20x/0.45 Ph1 objective (Nikon Corporation, Tokyo, Japan) and a Rolera XR 12 Bit Fast 1394 CCD camera (QImaging, Surrey, British Columbia) controlled by SimplePCI software (Hamamatsu). Excitation (340 and 380 nm, 50 ms exposure, mercury bulb) and emission light were separated using a 400 DCLP dichroic with a D510/80m filter. Data shown in Supplementary Figure 1 were acquired using a system comprised of an BX51WI (Olympus) microscope, a LUMPLFLN 40XW water-dipping lens (Olympus), a pE-340fura LED illumination system (CoolLED) and a SciCam Pro (Scientifica), controlled by μManager 2.0. Cells were perfused with HEPES-PSS using a gravity-fed perfusion system at a continuous rate of ~2 ml/min. Na^+^-free HEPES PSS was prepared as for HEPES-PSS above but with NaCl replaced with equimolar N-methyl-D-glucamine (0Na^+^/NMDG) or choline chloride (0Na^+^/ChoCl). Nominal Ca^2+^-free HEPES PSS was prepared as for HEPES-PSS above, omitting CaCl_2_.

### Drug preparation

Stocks of NS-1619, 2-APB, dantrolene, cyclopiazonic acid (CPA), and ionomycin were prepared in DMSO. The final DMSO concentration in working solutions was 0.1%. Stocks of La^3+^ were prepared using ultrapure water. Drugs were diluted into HEPES-PSS immediately prior to use.

### Measurement of [Ca^2+^]_i_ responses

Changes in [Ca^2+^]_i_ in response to treatment were measured as either the maximum fura-2 ratio change (ΔR_max_) or the area under the curve (AUC). The fura-2 fluorescence ratios for each trace were first normalised to the ratio at time 0 (R/R_0_) to control for variability in starting ratio between cells. To obtain a pretreatment baseline, the average ratio across the 6 timepoints (30 seconds at an acquisition interval of 5 seconds) immediately prior to treatment application was determined. ΔR_max_ was defined as the maximum increase from this baseline during the treatment period. The AUC was defined as the area under the curve during the entire treatment period (i.e. 10 min) in fura-2 ratio unit seconds (R.s). Where no [Ca^2+^]_i_ response was elicited in response to treatment, cells were stimulated with ionomycin (3 μM) to ensure a change in [Ca^2+^]_i_ could be measured. Cells in which no fura-2 fluorescence change was observed were excluded from analysis.

### Data analysis

Data analysis and statistical comparisons were performed using Microsoft Excel and GraphPad Prism 9. Statistical comparisons were performed using a nested t-test for side-by-side comparisons and a nested one-way ANOVA with post-hoc Tukey test for multiple comparisons. Within each figure, nested data are presented from individual experiments (3–6 independent repeats, each containing 12-40 independent cells) and the individual experimental means ± S.E.M.

## Results

### Na^+^ depletion drives an increase in [Ca^2+^]_i_ in breast cancer cells

Metastatic breast cancer cells exhibit elevated [Na^+^]_i_ (Leslie *et al*., 2019) that may regulate reverse-mode (Ca^2+^-entry) NCX activity (Pappalardo *et al*., 2014; Verkhratsky *et al*., 2018) in these cells. To assess the effects of lowering extracellular Na^+^ on [Ca^2+^]_i_, MDA-MB-231 cells were loaded with fura-2 AM (4 μM) and perfused with sequential pulses of HEPES-PSS where Na^+^ had been replaced with either equimolar N-methyl D glucamine (0Na^+^/NMDG) or equimolar choline chloride (0Na^+^/ChoCl) to remove extracellular Na^+^ while maintaining osmotic balance. Fura-2 fluorescence imaging revealed that, using either manoeuvre, removal of extracellular Na^+^ resulted in a steep increase in [Ca^2+^]_i_ that returned to baseline upon reperfusion with standard Na^+^-containing HEPES-PSS (Figures 1Ai and 1Bi). In parallel experiments, SBFI fluorescence imaging revealed that both 0Na^+^/NMDG and 0Na^+^/ChoCl resulted in a concomitant depletion of [Na^+^]_i_, suggesting a link between [Na^+^]_i_ depletion and subsequent [Ca^2+^]_i_. A similar increase in [Ca^2+^]_i_ upon perfusion with 0Na^+^/NMDG was observed in a second triple-negative breast cancer cell line, MDA-MB-468 (Supplementary Figure 1A). Interestingly, perfusion of 0Na/NMDG had no effect on [Ca^2+^]_i_ in the ER+ breast cancer cell line MCF-7 (Figure 1C), but induced a [Ca^2+^]_i_ transient in the Her2+ breast cancer cell line SKBR3 (Figure 1D); nevertheless, both MCF-7 and SKBR3 cells exhibited depletion of [Na^+^]_i_ upon perfusion with 0Na^+^/NMDG (Supplementary Figure 1B and 1C). Taken together, these data suggest that the [Ca^2+^]_i_ rise upon removal of extracellular Na^+^ and concomitant depletion of [Na^+^]_i_ is present only in certain breast cancer cell lines.

**Figure 1.**
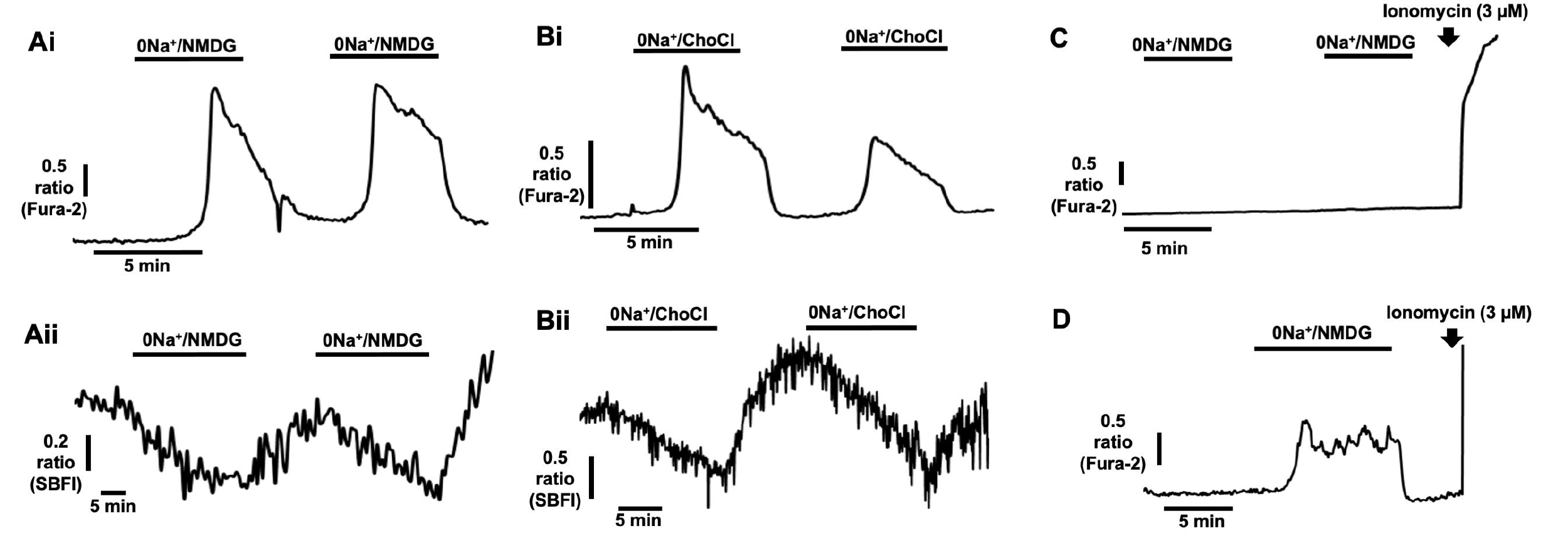
Na^+^-free conditions deplete [Na^+^]_i_ and induce elevated [Ca^2+^]_i_ in breast cancer cells. Fura-2 AM (4 μM) or SBFI AM (4 μM) fluorescence microscopy was used to measure [Ca^2+^]_i_ and [Na^+^]_i_ in cultured human breast cancer cells. Following perfusion with HEPES-PSS, MDA-MB-231 cells were perfused with Na^+^-free HEPES PSS; extracellular Na^+^ was replaced with equimolar N-methyl-D-glucamine (0Na^+^/NMDG) or choline chloride (0Na^+^/ChoCl) to maintain osmotic balance. Representative traces show the effects of 0Na^+^/NMDG and 0Na^+^/ChoCl on [Ca^2+^]_i_ (**Ai** and **Bi**, respectively) and [Na^+^]_i_ (**Aii** and **Bii**, respectively) in MDA-MB-231 cells. Additional experiments were performed in MCF-7 (**C**) and SKBR3 (**D**) cells. Ionomycin (3 μM) was used to elicit a [Ca^+^]_i_ increase as a positive control.

### The [Ca^2+^]_i_ increase induced by Na^+^-depletion is not due to Ca^2+^ influx from the extracellular space

We reasoned that the increase in [Ca^2+^]_i_ observed under 0Na^+^ conditions likely originated from one of two compartments: Ca^2+^ release from intracellular stores (most likely the endoplasmic reticulum) or Ca^2+^ influx from the extracellular space. To determine whether Ca^2+^ influx from the extracellular space contributed to the [Ca^2+^]_i_ responses observed, MDA-MB-231 cells were perfused with 0Na^+^/NMDG in nominal Ca^2+^-free conditions with La^3+^ (1 mM); La^3+^ in the millimolar range is a known inhibitor of both Ca^2+^ entry from and Ca^2+^ efflux to the extracellular space (James *et al*., 2013). La^3+^ was applied for 5 minutes prior to application of 0Na^+^/NMDG (Figure 2B). Compared with control cells (Figure 2A), La^3+^ had no effect on the [Ca^2+^]_i_ transients elicited following 0Na^+^/NMDG application, as measured by either the maximum change in fura-2 ratio (ΔRmax, Figure 2C) or the area under the curve (AUC, Figure 2D). These results indicate that Ca^2+^ influx via agonist-induced and store-operated Ca^2+^ entry mechanisms plays no role in the [Ca^2+^]_i_ transients induced following application of 0Na^+^/NMDG, and that the provenance of this Ca^2+^ response must be from some compartment other than the extracellular space.

**Figure 2.**
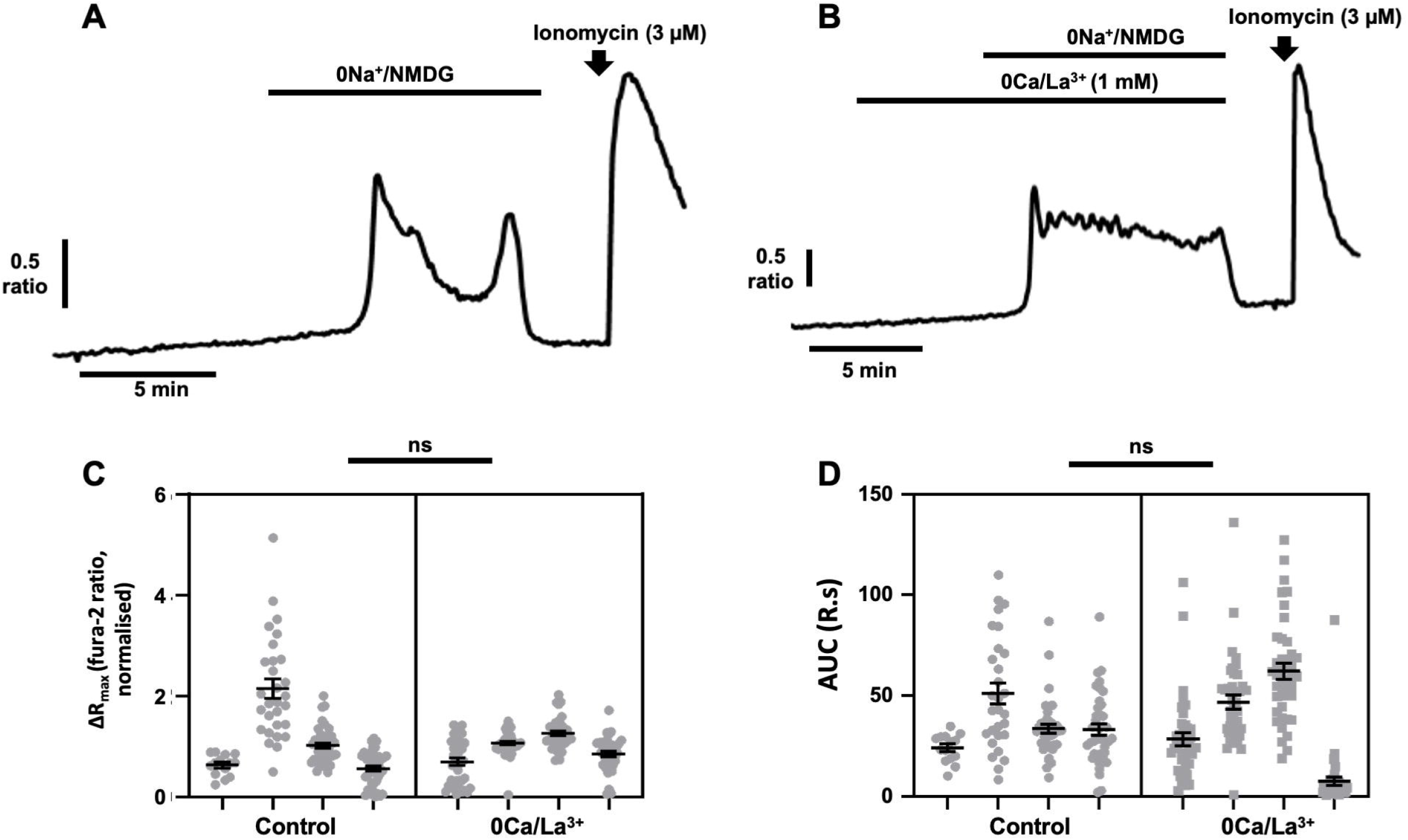
[Ca^2+^]_i_ transients induced by removal of extracellular Na^+^ in breast cancer cells are not due to Ca^2+^ entry. MDA-MB-231 cells were loaded with fura-2 AM (4 μM) and perfused with HEPES-PSS. Extracellular Na^+^ was replaced with equimolar 0Na^+^/NMDG to maintain osmotic balance (**A**). To block all Ca^2+^ entry from the extracellular space, cells were pretreated with five minutes of perfusion with Ca^2+^-free HEPES containing La^3+^ (1 mM, **B**). Ionomycin (3 μM) was applied at the end of an experiment to elicit a [Ca^+^]_i_ increase as a positive control. The maximum change in fluorescence ratio (ΔR_max_, **C**) and area under the curve (AUC, **D**) were compared using a nested t-test for side-by-side comparisons; n = 4 independent experiments for each condition, ns, not significant compared with control. Data presented are nested values for individual cells grouped by experiment (grey dots) and experimental means ± SEM (black line and bars).

### Elevations in [Ca^2+^]_i_ induced by Na^+^-depletion are not due to changes in membrane potential

Manoeuvres that deplete extracellular Na^+^ such as 0Na^+^/NMDG are known to cause V_m_ hyperpolarisation (Yang *et al*., 2020), thereby altering ion flux across the plasma membrane. Moreover, V_m_ hyperpolarization would increase the driving force for Ca^2+^ entry. To determine whether such changes in V_m_ led to the [Ca^2+^]_i_ transient observed under 0Na^+^/NMDG conditions, we applied the large-conductance Ca^2+^-activated K^+^ channel (K_Ca_1.1) activator NS-1619 at a concentration (10 μM) that hyperpolarises V_m_ in MDA-MB-231 cells to a similar degree to 0Na^+^/NMDG (Yang *et al*., 2020). Similar to control cells (Figure 3A), MDA-MB-231 cells treated with NS-1619 (10 μM, Figure 3B) showed no change in [Ca^2+^]_i_. However, an increased concentration of NS-1619 (40 μM, Figure 3C) elicited an increase in [Ca^2+^]_i_ that was significantly greater than any change observed in either the control cells (P<0.01 for both ΔRmax and AUC, Figures 3E and 3F) or those treated with 10 μM NS-1619 (P<0.01 for both ΔRmax and AUC, Figures 3E and 3F), indicating that dramatic changes in V_m_ can indeed alter [Ca^2+^]_i_. Importantly, the effects of 40 μM NS-1619 on [Ca^2+^]_i_ were abolished following pretreatment with La^3+^ (1 mM, P<0.05 for both ΔRmax and AUC, Figures 3C–F), indicating that the NS-1619 induced rise was due to Ca^2+^ influx from the extracellular space (Figure 3D). Given that Ca^2+^ influx plays no role in the [Ca^2+^]_i_ transients observed following application of 0Na^+^/NMDG (Figure 2), these results suggest that neither Ca^2+^ influx nor hyperpolarisation of V_m_ play a role in changes in [Ca^2+^]_i_ induced by application 0Na^+^/NMDG.

**Figure 3.**
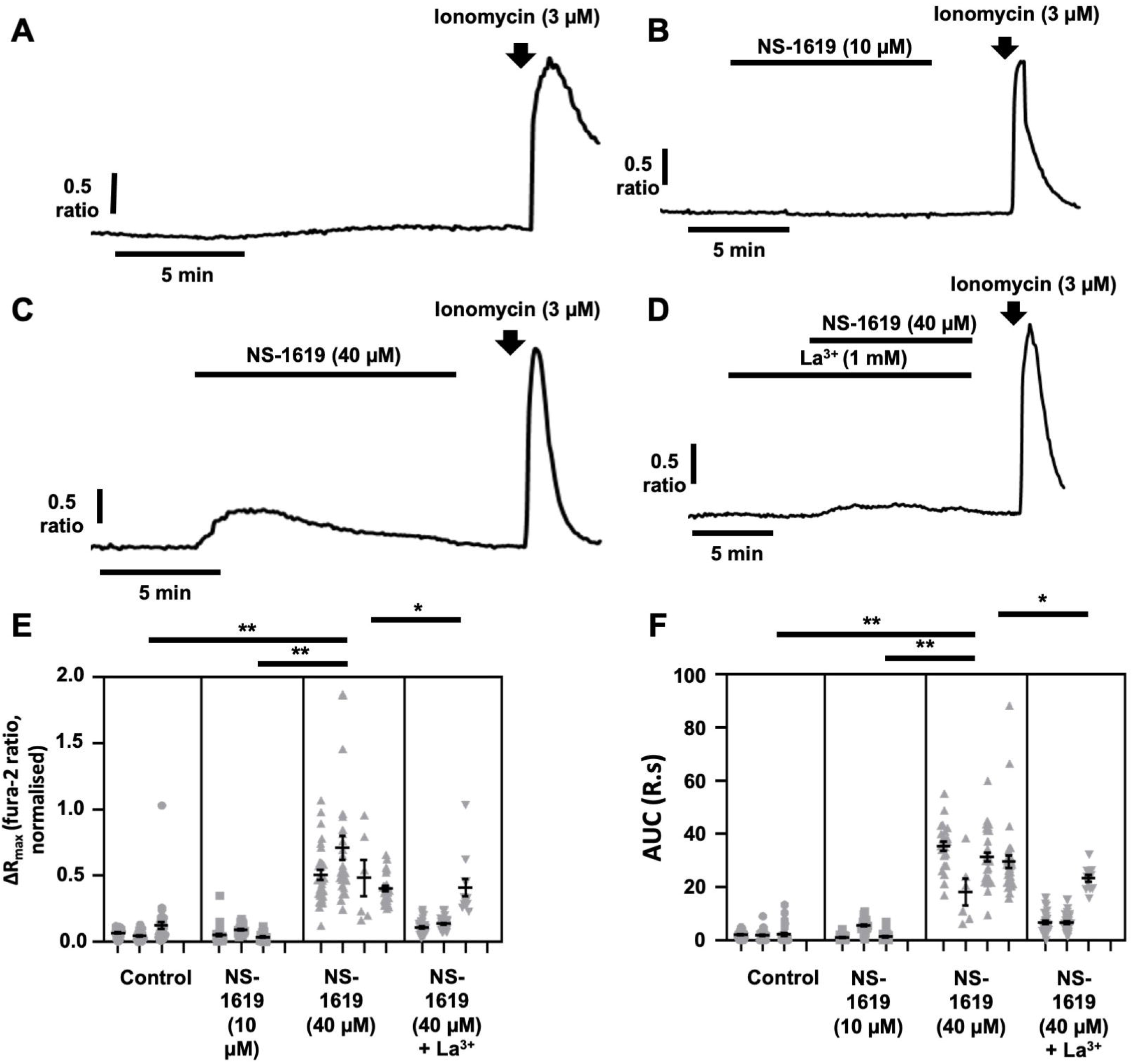
Hyperpolarisation of the membrane potential (V_m_) plays no role in [Ca^2+^]_i_ transients in breast cancer cells induced by removal of extracellular Na^+^. MDA-MB-231 cells were loaded with fura-2 AM (4 μM) and perfused with HEPES-PSS in the presence of the large-conductance Ca^2+^-activated K^+^ channel (K_Ca_1.1) activator NS-1619 to hyperpolarise V_m_. Representative traces are shown for cells perfused without NS-1619 (**A**) and with 10 μM NS-1619 (**B**) or 40 μM NS-1619 (**C**). Application of La^3+^ (**D**) determined that [Ca^2+^]_i_ changes observed following treatment with 40 μM NS-1619 were due to Ca^2+^ entry. Ionomycin (3 μM) was applied at the end of an experiment to elicit a [Ca^2+^]_i_ increase as a positive control. The maximum change in fluorescence ratio (ΔR_max_, **E**) and area under the curve (AUC, **F**) were compared using a nested one-way ANOVA with post-hoc Tukey test for multiple comparisons; n = 3–4 independent experiments for each condition. *, p < 0.05; **, p < 0.01 compared with control. Data presented are nested values for individual cells grouped by experiment (grey dots) and experimental means ± SEM (black line and bars).

### Depletion of ER Ca^2+^ stores inhibits the increase in [Ca^2+^]_i_ induced by 0Na^+^/NMDG

Having ruled out Ca^2+^ influx as being responsible for the [Ca^2+^]_i_ transients induced following application of 0Na^+^/NMDG, we tested whether depletion of the intracellular Ca^2+^ stores within the endoplasmic reticulum affected the observed rises in [Ca^2+^]_i_. To deplete endoplasmic reticulum Ca^2+^ stores, cells were treated with the sarcoplasmic/endoplasmic reticulum Ca^2+^ ATPase (SERCA) inhibitor cyclopiazonic acid (CPA, 30 μM) in nominal Ca^2+^-free conditions (to prevent store-operated Ca^2+^ entry), resulting in a transient leak of Ca^2+^ from the endoplasmic reticulum that was subsequently cleared from the cell (Figure 4B), presumably by the plasma membrane Ca^2+^ ATPase (James *et al*., 2015). These experiments were performed without EGTA since it was observed that inclusion of EGTA (1 mM) within the Ca^2+^-free buffer inhibited the [Ca^2+^]_i_ increases elicited by 0Na^+^/NMDG (Supplementary Figure 2). Compared with control cells, (Figure 4A), pretreatment with CPA completely abolished the increase in [Ca^2+^]_i_ elicited by 0Na^+^/NMDG (Figure 4B, 4C and 4D; P<0.01 for both ΔRmax and AUC). These results indicate that the 0Na^+^/NMDG-induced [Ca^2+^]_i_ transient is likely due to release of Ca^2+^ from the intracellular endoplasmic reticulum Ca^2+^ stores.

**Figure 4.**
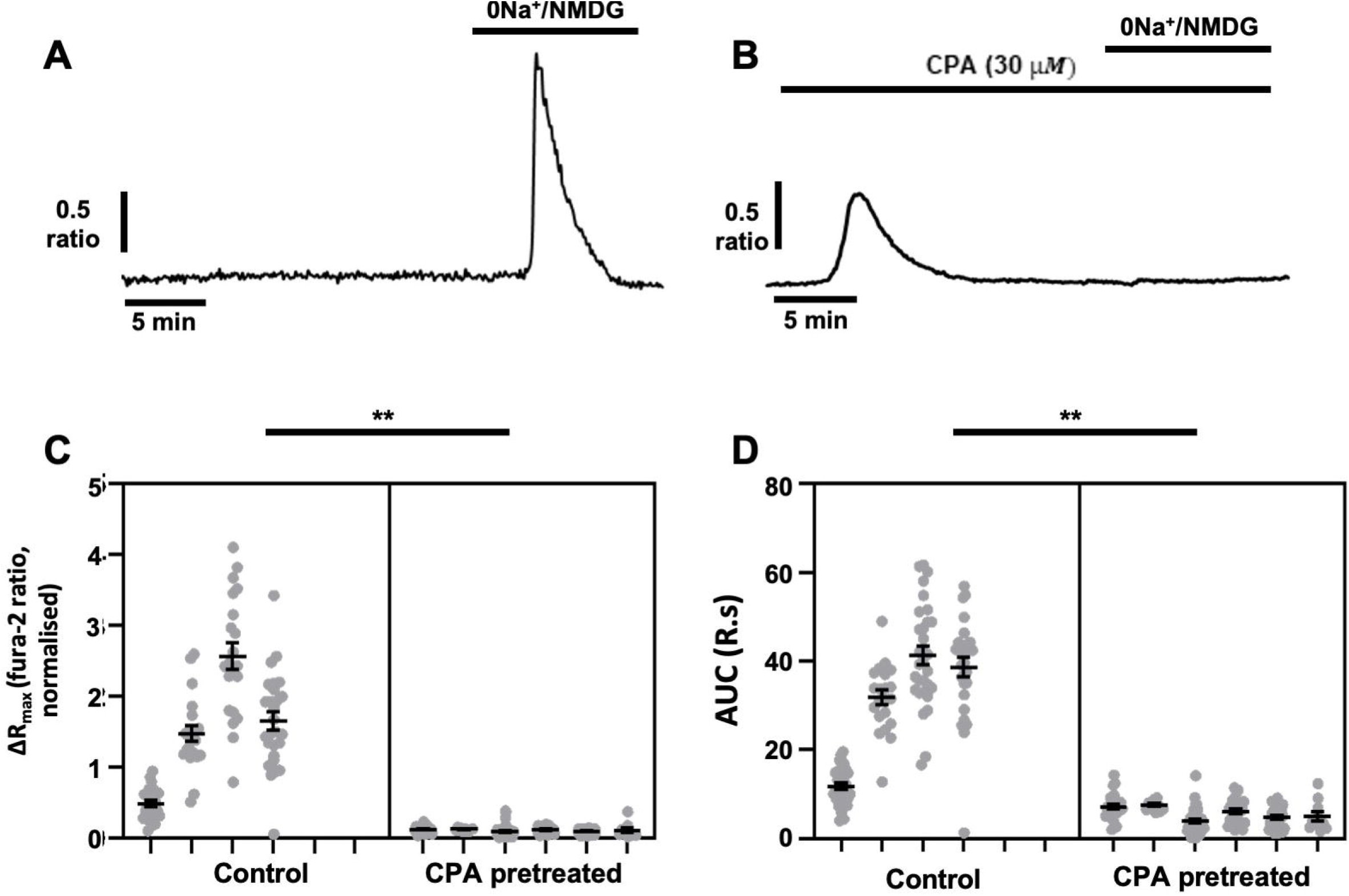
[Ca^2+^]_i_ transients induced by removal of extracellular Na^+^ in breast cancer cells require ER Ca^2+^ stores. MDA-MB-231 cells were loaded with fura-2 AM (4 μM) and Ca^2+^ imaging performed during replacement of extracellular Na^+^ with equimolar (0Na^+^/NMDG) in the presence or absence of the SERCA inhibitor cyclopiazonic acid (CPA, 30 μM). Representative traces are shown for cells perfused without CPA (**A**) or where cells were pretreated for 20 minutes CPA prior to application of 0Na^+^/NMDG (**B**). The maximum change in fluorescence ratio (ΔR_max_, **C**) and area under the curve (AUC, **D**) were compared using a nested t-test for side-by-side comparisons; n = 4–6 independent experiments for each condition; **, p < 0.01 compared with control. Data presented are nested values for individual cells grouped by experiment (grey circles) and experimental means ± SEM (black line and bars)

### Na^+^ depletion drives increases in [Ca^2+^]_i_ via a G-protein coupled receptor/IP_3_ receptor-mediated mechanism

Given the evidence implicating endoplasmic reticulum Ca^2+^ release in the changes in [Ca^2+^]_i_ observed upon application of 0Na^+^/NMDG, we next sought to determine the key molecular players responsible. Ca^2+^ is classically released from the endoplasmic reticulum via inositol (1,4,5) triphosphate receptors (IP_3_Rs) and ryanodine receptors (RyRs) (Berridge *et al*., 2000). To determine which of these release mechanisms was responsible for [Ca^2+^]_i_ transients upon application of 0Na^+^/NMDG, we perfused MDA-MB-231 cells with 0Na^+^/NMDG following pretreatment (10 min) with either the IP_3_R inhibitor 2-APB (50 μM) or the RyR inhibitor dantrolene (10 μM). Compared to control cells (Figures 5Ai and 5Bi), 2-APB significantly reduced the [Ca^2+^]_i_ transient induced by 0Na^+/^NMDG (Figures 5Aii, Aiii and Aiv; ΔRmax, P<0.05; AUC, P<0.001), whereas dantrolene (10 μM, Figure 5Bii) had no effect (Figure 5Biii and 5Biv). These data suggest that Na^+^ depletion induces Ca^2+^ release from the endoplasmic reticulum via IP_3_Rs.

**Figure 5.**
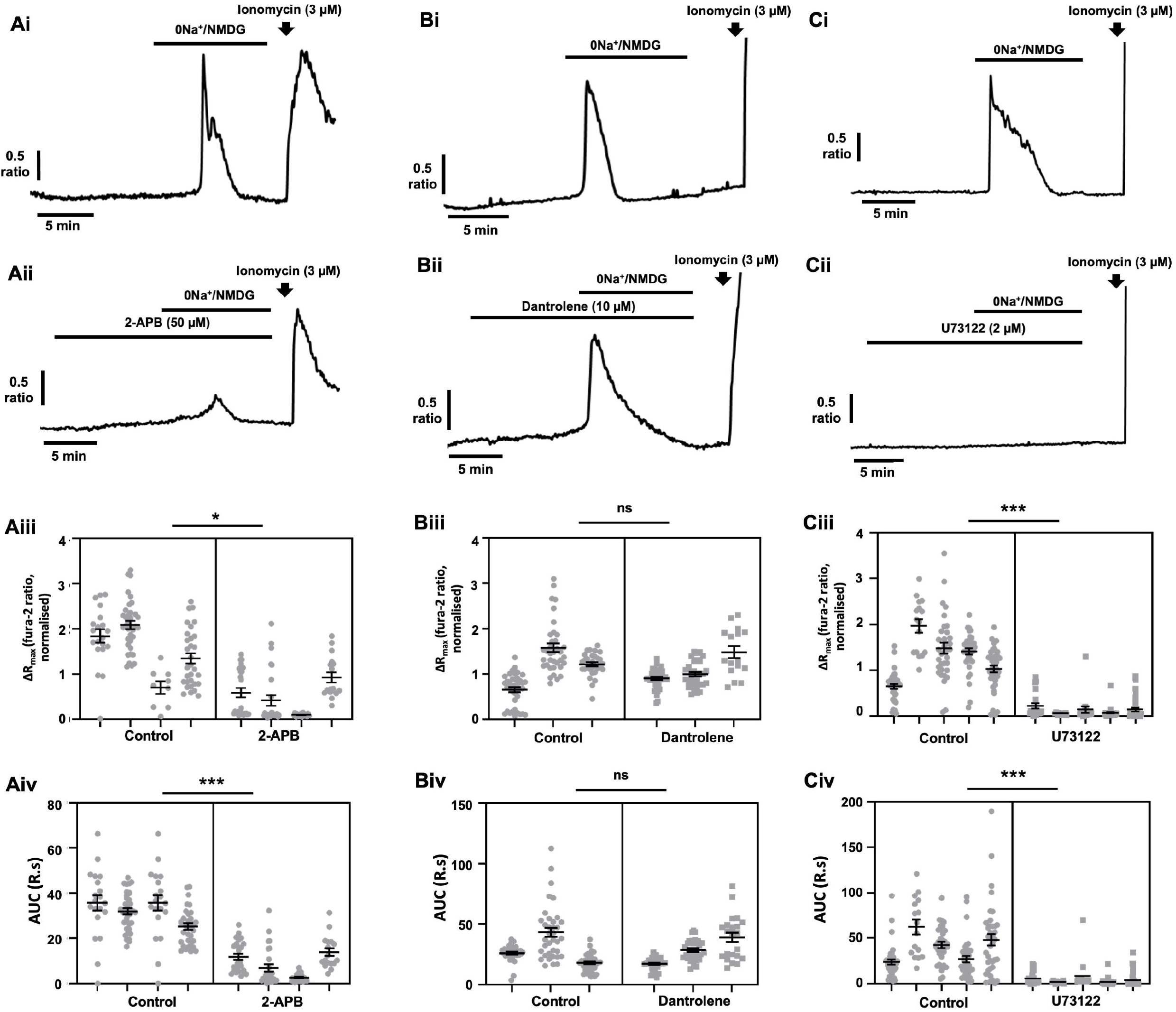
[Ca^2+^]_i_ transients induced by removal of extracellular Na^+^ require activation of IP_3_ receptor and phospholipase C, but are independent of ryanodine receptor activation. MDA-MB-231 cells were loaded with fura-2 AM (4 μM) and Ca^2+^ imaging performed during replacement of extracellular Na^+^ with equimolar (0Na^+^/NMDG) in the absence or presence of the IP_3_ receptor inhibitor 2-APB (50 μM, **Ai** and **Aii,** respectively), the ryanodine receptor inhibitor dantrolene (10 μM, **Bi** and **Bii**, respectively), and the phospholipase C inhibitor U73122 (2 μM, **Ci** and **Cii**, respectively). For each condition, cells were pretreated with drug for 10 minutes prior to application of 0Na^+^/NMDG. lonomycin (3 μM) was applied at the end of an experiment to elicit a [Ca^+^]_i_ increase as a positive control. The maximum change in fluorescence ratio (ΔR_max_; **Aiii**, 2-APB; **Biii**, dantrolene; **Ciii**, U73122) and area under the curve (AUC**; Aiv**, 2-APB; **Biv**, dantrolene; **Civ**, U73122) were compared using a nested t-test for side-by-side comparisons; n = 3–5 independent experiments for each condition; *, p < 0.05; ***, p < 0.001 compared with control. Data presented are nested values for individual cells grouped by experiment (grey dots) and experimental means ± SEM (black line and bars).

IP_3_Rs are classically activated by upstream G-protein coupled receptor signalling, with GPCRs of the G_αq_/_11_ family activating phospholipase C (PLC), which then cleaves membrane-bound phosphatidylinositol 4,5-bisphosphate (PIP2) to form the second messengers IP_3_ and diacylglycerol (DAG) (Patterson *et al*., 2005). To determine whether application of 0Na^+^/NMDG influenced IP_3_Rs via GPCR-mediated activation of PLC, we applied 0Na^+^/NMDG following pretreatment with the aminosteroid PLC inhibitor U73122 (2 μM, (Yule & Williams, 1992)). Compared with control cells (Figure 5Ci), U73122 completely abolished the increase in [Ca^2+^]_i_ observed following application of 0Na^+^/NMDG (Figure 5Cii, Ciii and Civ; P<0.001 for both ΔRmax and AUC), indicating that PLC activation, potentially via GPCRs, is critical to this Na^+^-controlled ER Ca^2+^ release mechanism.

## Discussion

The present study is the first to describe a relationship in metastatic breast cancer cells between Na^+^ depletion and a subsequent dramatic rise in [Ca^2+^]_i_ that is mediated by a PLC- and IP_3_R-dependent mechanism. These results provide further evidence of an interplay between Ca^2+^ and Na^+^ signalling in breast cancer cells that may have important implications for understanding cancer cell physiology. Through pharmacological manoeuvres we rule out any role for reverse-mode NCX activity, V_m_ and Ca^2+^ entry in this response. We also found that CPA abolished the [Ca^2+^]_i_ transients elicited by 0Na^+^/NMDG application; CPA is a well characterised SERCA blocker, providing further indirect evidence that this phenomenon was mediated by an IP_3_R-mediated endoplasmic reticulum Ca^2+^ release mechanism. Furthermore, while [Ca^2+^]_i_ transients were observed upon Na^+^ depletion in MDA-MB-231, MDA-MB-468 and SKBR3 cells, no such response was observed in MCF-7 cells. This finding indicates that the Na^+^-dependent Ca^2+^ increase is not a universal feature across all breast cancer cell lines. Further work is required to determine the underlying reason for the difference in Na^+^ and Ca^2+^ signalling between different cancer cell lines.

The present study establishes that Na^+^ regulates a PLC/IP_3_-dependent Ca^2+^ release in metastatic breast cancer cells; however, the molecular identity of the Na^+^ ‘sensor’ that initiates this event remains unknown. Interestingly, Na^+^ has been described as an endogenous regulator of Class A GPCRs (White *et al*., 2018) via allosteric regulation of agonist binding (Katritch *et al*., 2014; Zarzycka *et al*., 2019; Agasid *et al*., 2021). While we cannot presently rule out whether it is the depletion of cytosolic [Na^+^] or removal of extracellular Na^+^ that induces this [Ca^2+^]_i_ transient in the current study, these previous studies provide a strong potential candidate for the underlying trigger mechanism. Indeed, in these studies, Na^+^ depletion appeared to increase basal GPCR activity in the absence of agonist, suggesting that Na^+^ may act as a negative modulator of GPCR activation (Katritch *et al*., 2014; Zarzycka *et al*., 2019; Agasid *et al*., 2021). Interestingly, recently obtained X-ray structures of Class A GPCRs have identified a Na^+^ binding site in the vicinity of the agonist binding site that is exposed to the extracellular space (Yuan *et al*., 2013; Katritch *et al*., 2014). Moreover, computer simulations have suggested Class A GPCRs exhibit a transient water-filled channel connecting this Na^+^ binding site to the cytoplasm (Yuan *et al*., 2014, 2015; Vickery *et al*., 2018; Hu *et al*., 2019). Na^+^ translocation via such a channel would be expected to be electrogenic, and thus impact upon V_m_ as well as Na^+^ handling (Katritch *et al*., 2014; Shalaeva *et al*., 2019). This has important implications for cancer cells exhibiting altered Na^+^ dynamics, since V_m_ is an important regulator of cancer cell behaviour (Yang & Brackenbury, 2022); moreover, the altered [Na^+^]_i_ observed in breast tumours relative to healthy tissue (James *et al*., 2022) might be expected to disrupt GPCR regulation by Na^+^.

It is worthy to note that, in the present study, the PLC inhibitor U73122 had a more potent inhibitory effect on the [Ca^2+^]_i_ rise than the IP_3_R inhibitor 2-APB; the reasons for this are not clear, and suggest that U73122 is a more effective inhibitor of this pathway than 2-APB. However, 2-APB has been shown to target other channels, including certain TRP channel subtypes (Singh *et al*., 2018), which may have contributed to altered cation concentrations during these experiments. These findings also present a cautionary tale for cell physiologists using the methods employed in this study for exploring the relationship between Na^+^ conductance, Ca^2+^ signalling and wider cell behaviour. Application of 0Na^+^/NMDG or 0Na^+^/ChoCl is widely utilised for isolating cell responses from the influence of extracellular Na^+^; however, these manoeuvres are not inert (Thuma & Hooper, 2018). In addition to assessing for reverse mode NCX activity (Chovancova *et al*., 2020), Na^+^ regulation of GPCRs (and resultant effects on Ca^2+^ dynamics) should be taken into careful consideration and controlled for when using 0Na^+^/NMDG, 0Na^+^/ChoCl or similar manoeuvres to probe Na^+^ conductance mechanisms.

Interestingly, inclusion of EGTA in the Ca^2+^-free extracellular buffer abolished the [Ca^2+^]_i_ transients elicited by 0Na^+^/NMDG application, suggesting that EGTA exerts an inhibitory effect on the PLC and IP3 mediated Ca^2+^ release upon Na^+^ depletion. It has previously been shown that EGTA can inhibit mobilisation of IP_3_-sensitive Ca^2+^ stores (Finch & Goldin, 1994; Combettes & Champeil, 1994) in cerebellar microsomes. However, EGTA is considered membrane impermeable, and thus would not be expected to exert a direct effect on IP_3_Rs in the present study. This suggests that EGTA instead exerts its effects at the extracellular side of the plasma membrane via an at present undefined mechanism. Such a mechanism could be either by a direct effect on the putative GPCR involved or via chelation of extracellular divalent cations required for proper transmembrane protein function. Alternatively, it has been argued that extracellularly-applied EGTA may be taken up into cells via highly Ca^2+^-permeable cation channels (e.g. TRPA1, TRPV1, P2X7) and/or via fluid-phase endocytosis and subsequent release into the cytosol (Liu *et al*., 2020). Further work is required to resolve these possibilities.

Beyond the present study, very little is known about the link between Ca^2+^ signalling and Na^+^ homeostasis in the context of breast cancer. Evidence indicates that Ca^2+^ oscillations in MDA-MB-231 cells are regulated by aberrantly expressed VGSCs (Rizaner *et al*., 2016). Moreover, elevations in extracellular Na^+^ cause p-glycoprotein-induced resistance to paclitaxel in breast cancer cells via Ca^2+^ signalling (Babaer *et al*., 2018). Of particular relevance to the current study (where Na^+^ depletion activated IP_3_R-dependent Ca^2+^ signalling), IP3Rs are upregulated in breast cancer (Foulon *et al*., 2022), and estradiol-induced IP3R signalling regulates breast cancer cell proliferation (Szatkowski *et al*., 2010). Either pharmacological inhibition or knockdown of IP_3_Rs in breast cancer cells led to dysregulated bioenergetics, and autophagy, cell cycle arrest and cell death (Mound *et al*., 2013; Singh *et al*., 2017). Thus, the emerging evidence for altered Na^+^ and Ca^2+^ handling and their inextricable link via common channels and transporters hint at a hitherto relatively underexplored avenue in the context of cancer. The present study provides further evidence of such a link between Ca^2+^ and Na^+^ signalling that may have important implications for regulating cancer cell behaviour and future therapy.

## Author contributions

The project was designed by WJB and ADJ; experiments were carried out by ADJ, NS, KU, IJ and RS with the assistance of GJOE; data analysis was performed by ADJ, NS, KU, IJ and RS; the manuscript was prepared by ADJ and WJB; ADJ, WJB, GJOE and SC contributed to study design, interpretation of the data and critical revision of the paper for important intellectual content.

## Acknowledgements

This work was supported by Cancer Research UK (A25922) and an EPSRC Impact Accelerator Award to WJB and a Royal Thai Government Scholarship to NS.

## Disclosure statement

The authors declare that they have no competing interests.

## Notes

### Competing Interest Statement

The authors have declared no competing interest.

### Summary of Updates

Updates to methods and discussion.

