## Supplementary Figure 1 for "Sodium regulates PLC and IP_3_R-mediated calcium signalling in invasive breast cancer cells"

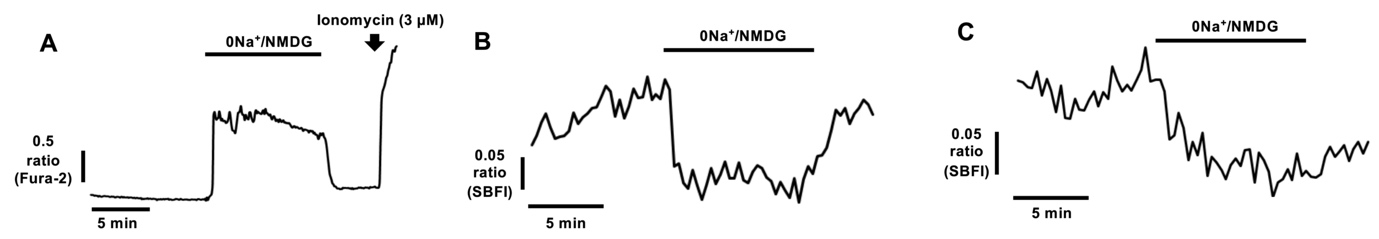

**Supplementary Figure 1: Na^+^-free conditions induce a [Ca^2+^]_i_ rise in MDA-MB-468 cells and deplete [Na^+^]_i_ in MCF-7 and SKBR3 cells.** Fura-2 AM (4 µM) or SBFI AM (4 µM) fluorescence microscopy was used to measure [Ca^2+^]_i_ and [Na^+^]_i_ in cultured human breast cancer cells. Following perfusion with HEPES-PSS, cells were perfused with Na^+^-free HEPES PSS; extracellular Na^+^ was replaced with equimolar N-methyl-D-glucamine (0Na^+^/NMDG) to maintain osmotic balance. Representative traces show the effects of 0Na^+^/NMDG on [Ca^2+^]_i_ in MDA-MB-468 cells (A) and on [Na^+^]_i_ in MCF-7 (B) and SKBR3 (C) cells.
