## Supplementary Figure 2 for "Sodium regulates PLC and IP_3_R-mediated calcium signalling in invasive breast cancer cells"

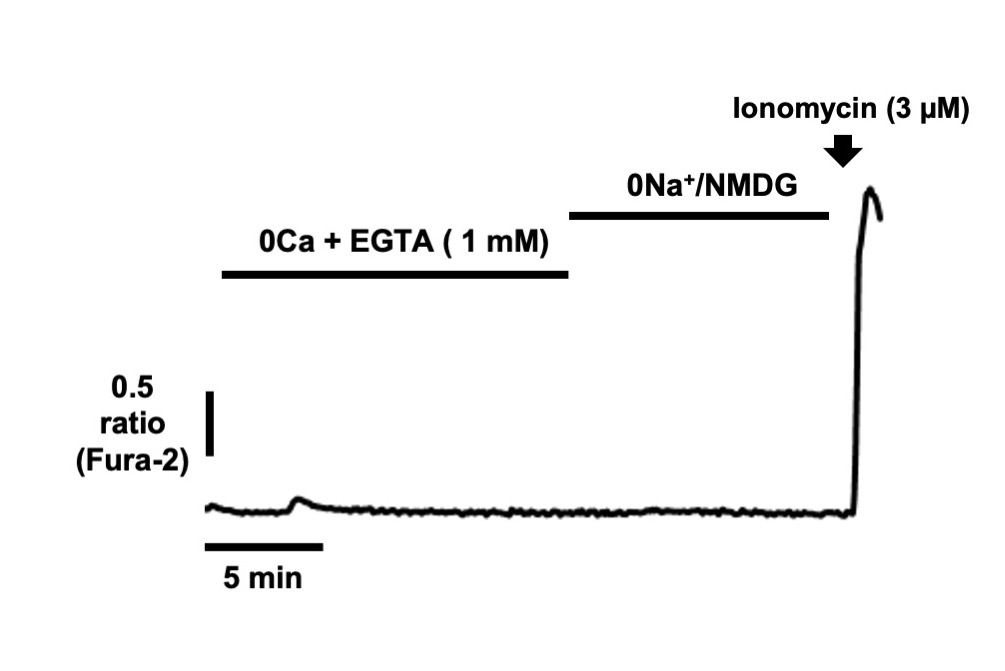

**Supplementary Figure 2: EGTA inhibits [Ca^2+^]_i_ transients induced by removal of extracellular Na^+^.** MDA-MB-231 cells were loaded with fura-2 AM (4 µM) and Ca^2+^ imaging performed during removal of extracellular Ca^2+^ in the presence of EGTA (1 mM), followed by replacement of extracellular Na^+^ with equimolar (0Na^+^/NMDG). Ionomycin (3 µM) was applied at the end of an experiment to elicit a [Ca^+^]_i_ increase as a positive control.
